# Cerebellar and cortico-striatal-midbrain contributions to reward-cognition processes and apathy within the psychosis continuum

**DOI:** 10.1101/2022.02.09.479617

**Authors:** Indrit Bègue, Janis Brakowski, Erich Seifritz, Alain Dagher, Philippe N. Tobler, Matthias Kirschner, Stefan Kaiser

## Abstract

Negative symptoms in the psychosis continuum are linked to impairments in reward processing and cognitive function. Processes at the interface of reward processing and cognition and their relation to negative symptoms remain little studied, despite evidence suggestive of integration in mechanisms and neural circuitry. Here, we investigated brain activation during reward-dependent modulation of working memory (WM) and their relationship to negative symptoms in subclinical and early stages of the psychosis continuum. We included 27 persons with high schizotypal personality traits and 23 patients with first episode psychosis as well as 27 healthy controls. Participants underwent functional magnetic resonance imaging while performing an established 2-back WM task with two reward levels (5 CHF vs. no reward), which allowed us to assess common reward-cognition regions through whole-brain conjunction analyses and to investigate relations with clinical scores of negative symptoms. As expected for behavior, reward facilitated performance while cognitive load diminished it. At the neural level, the conjunction of high reward and high cognitive load contrasts across the psychosis continuum showed increased hemodynamic activity in the thalamus and the cerebellar vermis. During high cognitive load, more severe apathy but not diminished expression in the psychosis continuum was associated with reduced activity in right lateral orbitofrontal cortex, midbrain, posterior vermal cerebellum, caudate and lateral parietal cortex. Our results suggest that hypoactivity in the cerebellar vermis and the cortical-striatal-midbrain-circuitry in the psychosis continuum relates to apathy possibly via impaired flexible cognitive resource allocation for effective goal pursuit.

## 1. Introduction

The psychosis continuum comprises a spectrum of psychopathological manifestations characterized by overlapping clinical and demographic risk factors as well as shared neuroimaging and genetic features(DeRosse and Karlsgodt 2015). It spans from schizotypy - a range of personality traits manifesting with social isolation, odd behavior and unusual perceptions(Campellone, Elis et al. 2016) – and early psychosis to chronic schizophrenia(van Os, Linscott et al. 2009, Nelson, Seal et al. 2013). Across the psychosis continuum, negative symptoms (NS) represent a core psychopathological domain that maps onto at least two dimensions: apathy and diminished expression(Kirkpatrick, Fenton et al. 2006, Sauvé, Brodeur et al. 2019, Begue, Kaiser et al. 2020). Apathy, in particular, has received increasing scientific and clinical attention in recent years due to its high impact on long-term disability and poor quality of life even in subclinical and early phases of psychosis(Milev, Ho et al. 2005, Cohen and Davis 2009, Gee, Hodgekins et al. 2016, Rammou, Fisher et al. 2019).

On a neurobiological level, emergence of apathy has been predominantly examined in schizophrenia and is conceptualized as resulting from impairments in goal-directed behavior with compelling evidence pointing to abnormal reward processing (reviewed in(Strauss, Waltz et al. 2014)). A second line of research has linked NS to deficits in executive function(Faerden, Vaskinn et al. 2009, Hartmann, Kluge et al. 2015, Raffard, Gutierrez et al. 2016) and in working memory (WM)(Barch 2003, Raffard, Gutierrez et al. 2016), in particular to impairments in WM resource mobilization to perform demanding cognitive tasks(Culbreth, Moran et al. 2020). Less attention has been dedicated to the convergence of reward and cognition processes in the psychosis continuum. Although ‘reward’ and ‘cognition’ are considered as distinct psychological constructs, they communicate in a bidirectional fashion and need to be integrated for goal-directed behavior. In fact, motivated behavior requires internal representation and active maintenance of cognitive goals in neural hubs(Pessoa and Engelmann 2010, Braver, Krug et al. 2014, D’Esposito and Postle 2015) and expected rewards impact how information pertaining to cognitive function is represented(Krawczyk, Gazzaley et al. 2007, Kennerley and Wallis 2009). Importantly, allocation of WM resources *per se*, as an effortful process, requires motivation-related dopaminergic input to transform motivation-enhancing incentives into cognitive performance (reviewed in(Westbrook and Braver 2016)). From a neural perspective, goal-directed behavior relies on integration of fronto-parietal, fronto-striatal and cortico-thalamo-cerebellar neural circuitries(Brown and Pluck 2000, Pessoa 2009, Braver, Krug et al. 2014, Caligiore, Pezzulo et al. 2017, Parro, Dixon et al. 2018).

Here, we integrate these two separate lines of research in one common experimental setting using an established incentivized 2-back task(Pochon, Levy et al. 2002) focusing on early stages of the psychosis continuum. Our aim is to examine at the neural level whether and how reward-cognition hubs play a role in apathy pathophysiology across the psychosis continuum. Capitalizing on a sample that includes unmedicated individuals with high schizotypy and minimally medicated first-episode patients, we 1) adopt a whole brain approach to allow examining neural regions beyond classic mesolimbic correlates of negative symptoms 2) delineate brain regions implicated not only in the primary processes (e.g. reward or cognitive function) but positioned at their intersection, in other words, shared reward-cognitive neural signatures as identified by a conjunction analysis. Finally, we sought to 3) examine dimensional associations with negative symptom subdomains across clinical and sub-clinical stages of psychosis.

## 2. Methods

### 2.1. Participants

We assessed 52 individuals of the psychosis continuum, namely 27 healthy individuals with high schizotypy (SPT), 25 patients with first episode non-affective psychosis (FEP) as well as 28 healthy control participants (HC), as a part of a larger study in the psychosis spectrum including schizophrenia(Hager, Kirschner et al. 2015, Kirschner, Hager et al. 2016, Kirschner, Haugg et al. 2018). We excluded two FEP and one HC participants due to excessive movements in the MRI. Thus, the final psychosis continuum sample (n=50) consisted of 27 healthy SPT individuals, 23 FEP patients and 27 healthy control participants. We excluded patients with major causes of secondary NS, i.e. with major depressive episode, significant extrapyramidal side effects (total score> 2 on the Modified Simpson–Angus Scale(Simpson and Angus 1970)), florid psychotic symptoms (any positive subscale item score 4 or higher on the Positive and Negative Syndrome Scale (PANSS)(Kay, Opler et al. 1987)) or excessive sedation (lorazepam>1 mg/d). Full details on participant recruitment and psychopathological assessment can be found in the Supplementary material.

### 2.2. MRI data Acquisition

Two runs containing 185 whole brain T2* weighted echo-planar images (EPI) using a Philips Achieva 3.0 T magnetic resonance scanner with a 32 channel SENSE head coil (Philips, Best, The Netherlands). Functional data acquisition parameters included: in-plane resolution of 3×3mm^2^, 3mm of slice thickness, 0.5mm gap width over a field of view of 240×240mm, repetition/echo time (TR/TE) of 2000/25ms, flip angle of 82°. Slices were acquired in ascending order and aligned with the anterior– posterior commissure. Structural T1-weighted image were acquired with the following parameters: 160 sagittal slices, with acquisition voxel size=1.0 × 1.0 × 1.0mm3, time between two inversion pulses=2997ms, inversion time = 1008 ms, inter-echo delay = 8.1 ms, flip angle = 8°, matrix = 240 × 240, field of view =240 × 240 mm2.

### 2.3. fMRI Task

We used a modified version of a previously employed letter n-back task applied in healthy volunteers and patients with schizophrenia(Pochon, Levy et al. 2002, Owen, McMillan et al. 2005, Hager, Kirschner et al. 2015), see Figure 1 for details.

**Figure 1.**
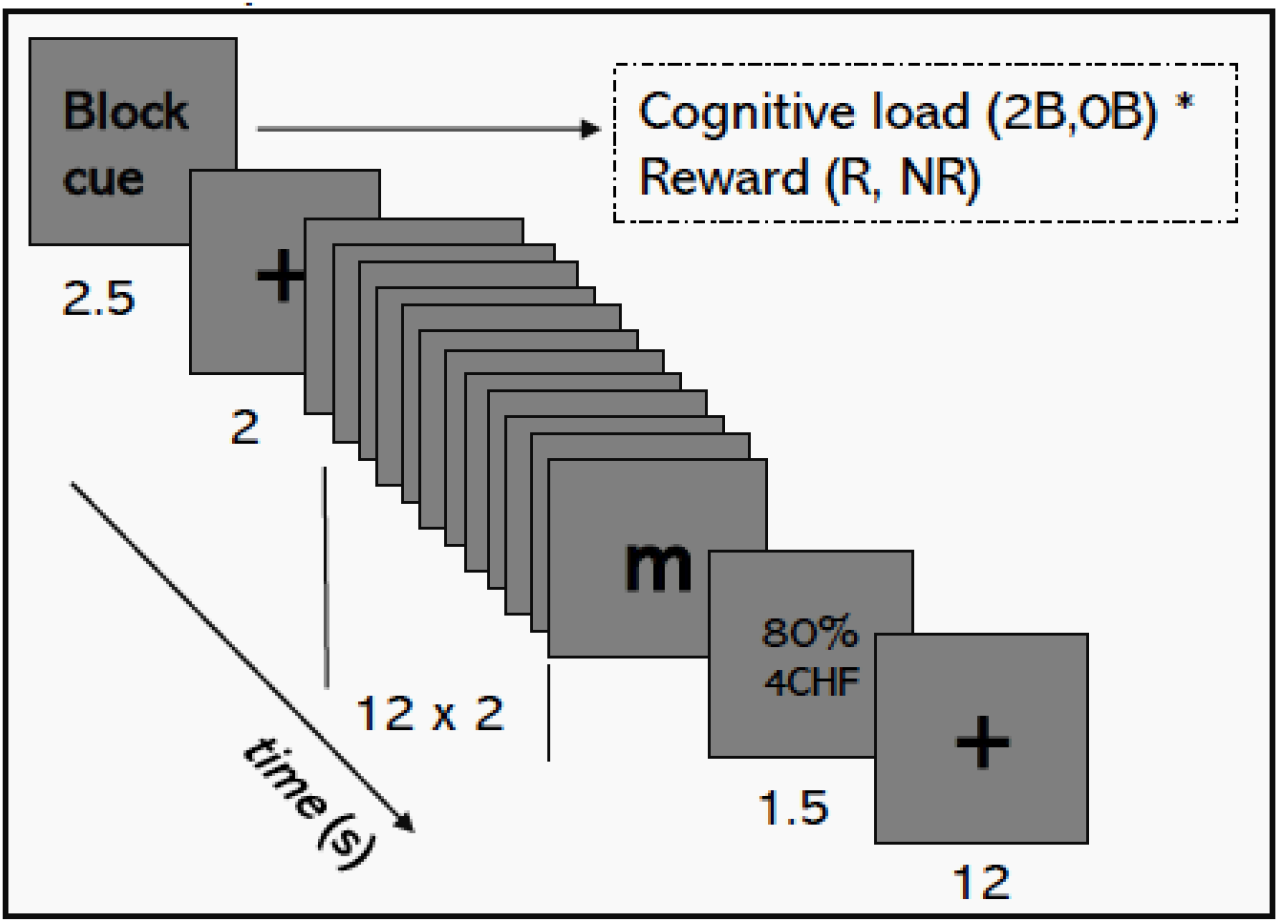
Schematic representation of the modified letter n-back task. The task was constructed in a 2×2 factorial design [cognitive load (0-back vs. 2-back) and reward (reward vs. no reward)], resulting in four different conditions: 0-back/reward (0R), 0-back/no reward (0N), 2-back/reward (2R), 2-back/no reward (2N), with 4 blocks per condition split in 2 runs. The order of presentation was fixed for all subjects and as follows: 0R, 2R, 0N, 2N, 2N, 0N, 2R, 0R; 0R, 0N, 2R, 2N, 2N, 2R, 0N, 0R. Participants had to press a button whenever the letter *x* letter appeared on the screen (0-back condition), or whenever the letter on the screen was identical to 2 letters ago (2-back task). Participants received monetary reward according to their performance (maximum payment per block was 5 Swiss Francs (CHF) and minimum payment was 0 CHF) in the rewarded conditions and no payment in the unrewarded conditions. Participants received a guaranteed amount of 10 CHF. After indication of the current condition, a fixation cross followed. One block consisted of 12 letter stimuli containing 4 targets. Each letter appeared for 500 ms and was followed by an inter-stimulus interval of 1500 ms. After the presentation of all 12 stimuli, a feedback about the performance and the monetary gain was given for 1500 ms. A resting period of 12,000 ms followed after every block. Abbreviations: 0B=0-back task; 2B=2-back task; NR= no reward; R=reward; HC=healthy controls;

### 2.4. Behavioral Data Analyses

We analyzed demographic, clinical and neuropsychological data using Statistica® (Statistica 8, Statsoft, Tulsa, USA). Primary behavioral outcome measure of our n-back task was the sensitivity index d’(Haatveit, Sundet et al. 2010) defined as the standardized probability of a hit minus the standardized probability of a false alarm. As an additional outcome parameter, we measured the response time (RT) of correct trials in each condition. To test for differences in behavioral performance, d’ and response times were entered into separate mixed-design ANOVAs with group (HC, psychosis continuum) as between-subjects factor and cognitive load [0-back (0B), 2-back (2B)] and reward [no reward (N), reward (R)] as within-subject factors.

### 2.5. Image Analyses

#### 2.5.1. Preprocessing

We used a standard pipeline using SPM12 (Statistical Parametric Mapping, Wellcome Department of Cognitive Neurology, London, UK) including 1) realignment 2) co-registration of the structural scan to the mean EPI images 3) segmentation of the structural image and subsequent application of the issuing estimated deformation field to the functional images, 4) normalization into standard stereotaxic space and resampling at 3×3×3 mm3 voxel size 5) smoothing of the normalized images with a 6 mm full-width at half-maximum isotropic Gaussian kernel. The quality of fMRI data was assessed by visual inspection (e.g. examining for significant signal dropout in echo planar image (EPI) sequences) and by a threshold of>3 mm for head movement.

#### 2.5.2. First level

For each group, we analyzed preprocessed images using the general linear model (GLM) framework implemented in SPM12 in a 2×2 design [2 levels of cognitive load (0-back and 2-back) and 2 levels of reward (reward and no reward)]. This design resulted in four experimental conditions: no cognitive load and no reward (0BN), no cognitive load and reward (0BR), cognitive load and no reward (2BN) and cognitive load and reward (2BR). For each condition, we convolved block epochs with the canonical hemodynamic response function using a box-car function with the respective block duration starting at the onset of each block. To account for head motion-related variance, we included the six differential parameters derived from the realignment process [x, y, and z translations (in millimeters) plus pitch, roll, and yaw rotations] as regressors of no interest. Low-frequency signal drifts were filtered using a cut-off period of 128 s. Global scaling was applied, with each fMRI value rescaled to a percentage value of the average whole-brain signal for that scan.

#### 2.5.3. Second level

We assessed contrast images corresponding to each condition of interest [cognitive load (2-back>0-back), reward (reward>no reward). These were fed as a within-subject factor into a flexible factorial analysis that modelled group by condition. The between-subject factor was group (HC, psychosis continuum (FEP + SPT)) and subject served as a random factor. We set each of these three factors to have unequal variance between their levels.

#### 2.5.4. Whole brain second level analyses

Given our focus here on shared regions of both reward and cognitive load processing, we performed a whole brain conjunction analysis (logical AND) for the contrasts of cognitive load and reward (2-back>0-back AND reward>no reward)(Price and Friston 1997). Conjunction analysis is particularly useful when examining consistent and common effects of two orthogonal contrasts(Krebs, Boehler et al. 2011, Soreq, Leech et al. 2019, Van Oudenhove, Kragel et al. 2020). To this end, we performed minimum T-statistic null conjunction analysis over 2 orthogonal contrasts(Friston, Penny et al. 2005). The null distribution for the minimum statistic is described by Friston and colleagues(Friston, Penny et al. 2005). This analysis enabled us to infer a conjunction of multiple effects at significant voxels. We then applied a threshold of p<.001 uncorrected with a whole-brain threshold of FWE p<.05 corrected at the cluster level to account for multiple comparisons and generate a corrected map of the conjunction effect.

We also asked whether there was stronger reward coding under high cognitive load than under low cognitive load at the neural level. Therefore, we computed the interaction contrast [(2-back reward – 2-back no reward)>(0-back reward – 0 back no reward)].

#### 2.5.5. Dimensional analyses

In order to establish brain-psychopathology relationships, we examined how regions active during cognitive load or reward processing and their interaction were differentially associated to negative symptom severity in the psychosis continuum. Therefore, we constructed a multiple linear regression model using the cognitive load contrast (2-back>0-back) with the BNSS apathy factor and BNSS diminished expression factor as linear regressors. We adjusted for depression effects by including in the model the individual CDS scores(Maillet, Krack et al. 2016). We performed the same regression analysis for the reward contrast (reward>no reward) and the interaction contrast [(2-back reward – 2-back no reward)>(0-back reward – 0 back no reward)]. Please note that only one conjunction map can be computed at the population level, thus such regression mapping of subject-level effects to individual scores cannot be performed for the conjunction analysis.

Cluster extent was calculated based on p<.001 uncorrected and type I error was controlled by cluster correction for multiple voxel comparison using a statistical threshold of FWE p<.05. Statistical maps are displayed using MRIcroGL (www.nitrc.org).

## 3. Results

### 3.1. Demographic, neuropsychological and psychopathology results

Demographic, clinical and neuropsychological data are summarized in Table 1. A summary of the relationship between negative symptom dimensions (apathy and diminished expression) and depression, general and psychosocial functioning, positive symptoms, cognitive function and medication dosage is represented in the Supplementary material (Table S1).

**Table 1.**
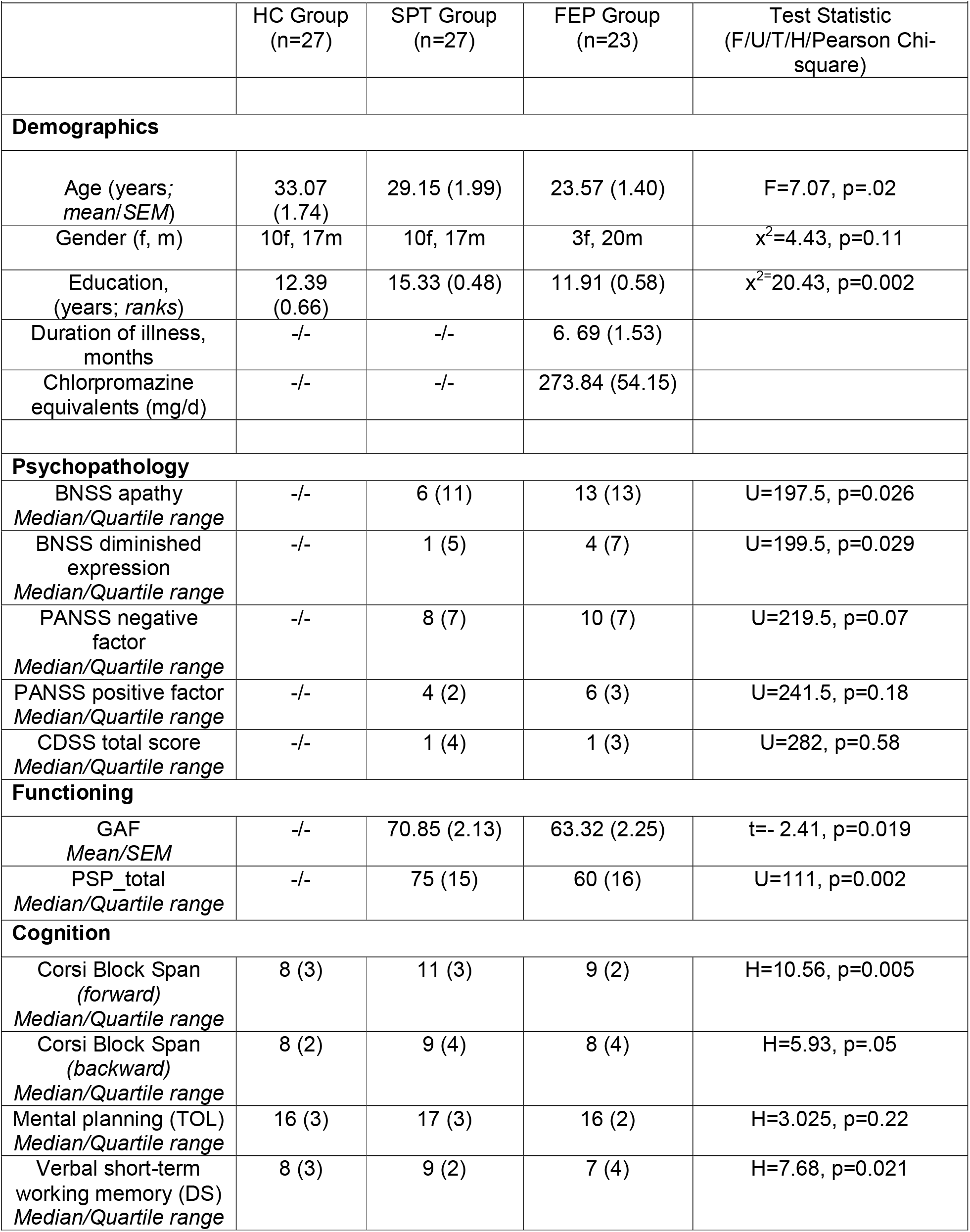
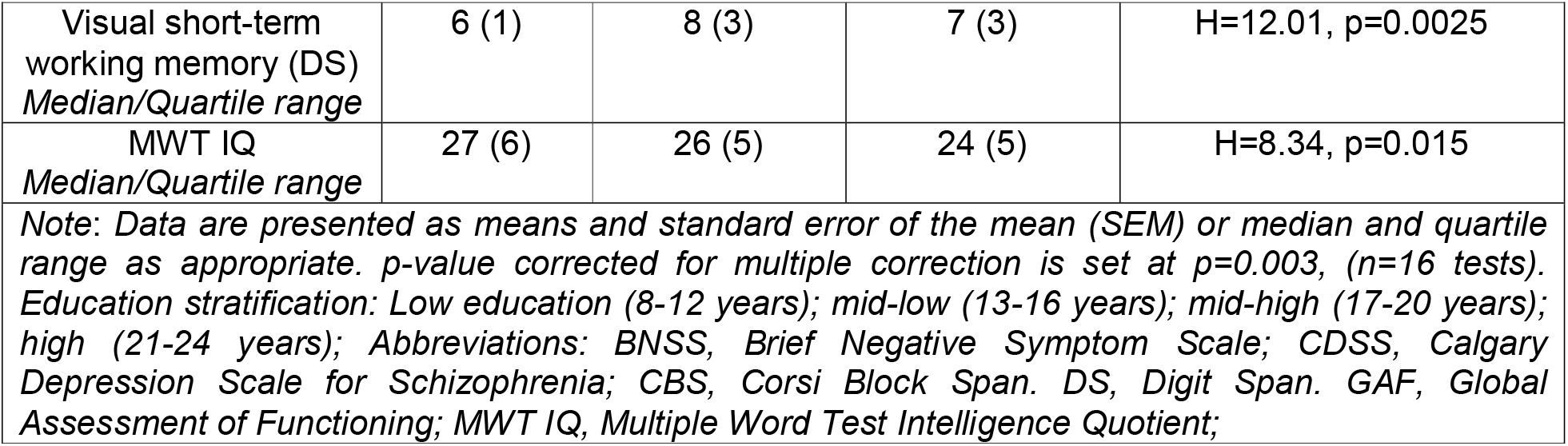
Summary of demographic characteristics, psychopathology and neurocognitive testing

|  |  |  |  |  |
| --- | --- | --- | --- | --- |
|  | HC Group<br>(n=27) | SPT Group<br>(n=27) | FEP Group<br>(n=23) | Test Statistic<br>(F/U/T/H/Pearson Chi-square) |
| **Demographics** |  |  |  |  |
| Age (years;<br>*mean/SEM*) | 33.07<br>(1.74) | 29.15 (1.99) | 23.57 (1.40) | F=7.07, p=.02 |
| Gender (f, m) | 10f, 17m | 10f, 17m | 3f, 20m | $\chi^2=4.43$, p=0.11 |
| Education,<br>(years; *ranks*) | 12.39<br>(0.66) | 15.33 (0.48) | 11.91 (0.58) | $\chi^2=20.43$, p=0.002 |
| Duration of illness,<br>months | -/- | -/- | 6. 69 (1.53) |  |
| Chlorpromazine<br>equivalents (mg/d) | -/- | -/- | 273.84 (54.15) |  |
| **Psychopathology** |  |  |  |  |
| BNSS apathy<br>*Median/Quartile range* | -/- | 6 (11) | 13 (13) | U=197.5, p=0.026 |
| BNSS diminished<br>expression<br>*Median/Quartile range* | -/- | 1 (5) | 4 (7) | U=199.5, p=0.029 |
| PANSS negative<br>factor<br>*Median/Quartile range* | -/- | 8 (7) | 10 (7) | U=219.5, p=0.07 |
| PANSS positive factor<br>*Median/Quartile range* | -/- | 4 (2) | 6 (3) | U=241.5, p=0.18 |
| CDSS total score<br>*Median/Quartile range* | -/- | 1 (4) | 1 (3) | U=282, p=0.58 |
| **Functioning** |  |  |  |  |
| GAF<br>*Mean/SEM* | -/- | 70.85 (2.13) | 63.32 (2.25) | t=- 2.41, p=0.019 |
| PSP_total<br>*Median/Quartile range* | -/- | 75 (15) | 60 (16) | U=111, p=0.002 |
| **Cognition** |  |  |  |  |
| Corsi Block Span<br>(forward)<br>*Median/Quartile range* | 8 (3) | 11 (3) | 9 (2) | H=10.56, p=0.005 |
| Corsi Block Span<br>(backward)<br>*Median/Quartile range* | 8 (2) | 9 (4) | 8 (4) | H=5.93, p=.05 |
| Mental planning (TOL)<br>*Median/Quartile range* | 16 (3) | 17 (3) | 16 (2) | H=3.025, p=0.22 |
| Verbal short-term<br>working memory (DS)<br>*Median/Quartile range* | 8 (3) | 9 (2) | 7 (4) | H=7.68, p=0.021 |
| Visual short-term working memory (DS)<br>*Median/Quartile range* | 6 (1) | 8 (3) | 7 (3) | H=12.01, p=0.0025 |
| MWT IQ<br>*Median/Quartile range* | 27 (6) | 26 (5) | 24 (5) | H=8.34, p=0.015 |
| *Note: Data are presented as means and standard error of the mean (SEM) or median and quartile range as appropriate. p-value corrected for multiple correction is set at p=0.003, (n=16 tests). Education stratification: Low education (8-12 years); mid-low (13-16 years); mid-high (17-20 years); high (21-24 years); Abbreviations: BNSS, Brief Negative Symptom Scale; CDSS, Calgary Depression Scale for Schizophrenia; CBS, Corsi Block Span. DS, Digit Span. GAF, Global Assessment of Functioning; MWT IQ, Multiple Word Test Intelligence Quotient;* |  |  |  |  |

Overall, we found no significant correlation between apathy and positive symptom severity, medication dosage, or cognitive function. There was a negative association of apathy with global functioning and with psychosocial functioning. Despite low depression levels (Table 1), there was a significant association of apathy and depression (Table S1). Diminished expression correlated negatively with intelligence and verbal fluency (Table S1).

### 3.2. Behavioral results: effective experimental manipulations

Our behavioral outcome measures were sensitivity (d’) and response times during the 2-back task. Regarding sensitivity, we found a main effect of reward [F(1, 75)=7.68, p=0.007] and a main effect of cognitive load [F(1, 75)=8.8, p=0.004] but not a main effect of group (p=0.36) (Table 2). All interactions did not pass threshold for significance. Regarding response times, we found a main effect of reward [F(1, 75)=11.74, p=0.001] and a main effect of cognitive load [F(1,75)=49.86, p=0.71 x 10^-9^] but not a main effect of group (p=0.57) (Table 2). All interactions did not pass threshold for significance.

**Table 2.** Summary of behavioral results

| **Table 2.** Summary of behavioral results |  |  |
| --- | --- | --- |
|  | HC<br>(n=27)<br>mean, standard deviation | Psychosis continuum<br>(n=50)<br>mean, standard deviation |
| **Accuracy (d')** |  |  |
| 0-back reward | 7.12(0.80) | 7.20(0.51) |
| 0-back no reward | 6.98(0.92) | 7.17(0.52) |
| 2-back reward | 6.89(0.68) | 7.09(0.72) |
| 2 back no reward | 6.69(0.98) | 6.73(10.8) |
| **Response times (in ms)** |  |  |
| 0-back reward | 445.81(52.49) | 448.73(78.70) |
| 0-back no reward | 456.13(59.61) | 468.42(70.58) |
| 2-back reward | 511.01(106.30) | 518.78(108.92) |
| 2 back no reward | 524.83(114.86) | 544.80(123.57) |

**Table 3.** Summary of whole brain regression of cognitive load (2-back>0-back) and apathy across the psychosis continuum

| Region Label |  | Cluster extent | t-value | p<sub>FWE-CL</sub> | MNI Coordinates |  |  |
| --- | --- | --- | --- | --- | --- | --- | --- |
|  |  |  |  |  | x | y | z |
| R | Cerebellar Vermis | 57 | 4.26 | 0.016 | 6 | -60 | -30 |
| R | Caudate | 75 | 4.83 | 0.004 | 18 | 0 | 18 |
| R | Habenular complex | 161 | 5.67 | < .001 | 3 | -30 | 3 |
| R | Lateral OFC | 57 | 6.42 | 0.016 | 36 | 30 | -9 |
| R | Precuneus | 229 | 4.70 | < .001 | 3 | -63 | 39 |
| R | Middle Temporal Gyrus | 64 | 4.09 | 0.01 | 60 | -51 | 15 |
| L | Supramarginal | 103 | 5.13 | <.001 | -54 | -48 | 30 |
| R | Midbrain | 161 | 4.38 | < .001 | 3 | -30 | -27 |

Across all experimental conditions (0BR, 0BN, 2BR, 2BN), there was no specific association to either diminished expression or apathy for both indices of performance (d’, response times; none of the p-values passed the significance threshold) in the psychosis continuum. Thus, overall, our behavioral findings indicate that reward and cognitive load manipulations were effective in all groups and were not related to negative symptom dimensions.

### 3.3. Functional MRI results

#### 3.3.1. Effects of cognitive load

Cognitive load (2-back>0-back) yielded brain activity in the frontoparietal and cerebellar network (Table S2) in both the healthy controls and the psychosis continuum cohorts (Figure 2, A1 and B1). Cognitive load yielded more frontoparietal activity in the psychosis continuum versus healthy controls, whereas the opposite contrast revealed that cognitive load yielded more insular and mediofrontal activity in healthy controls versus patients in the psychosis continuum (see supplementary material).

**Figure 2.**
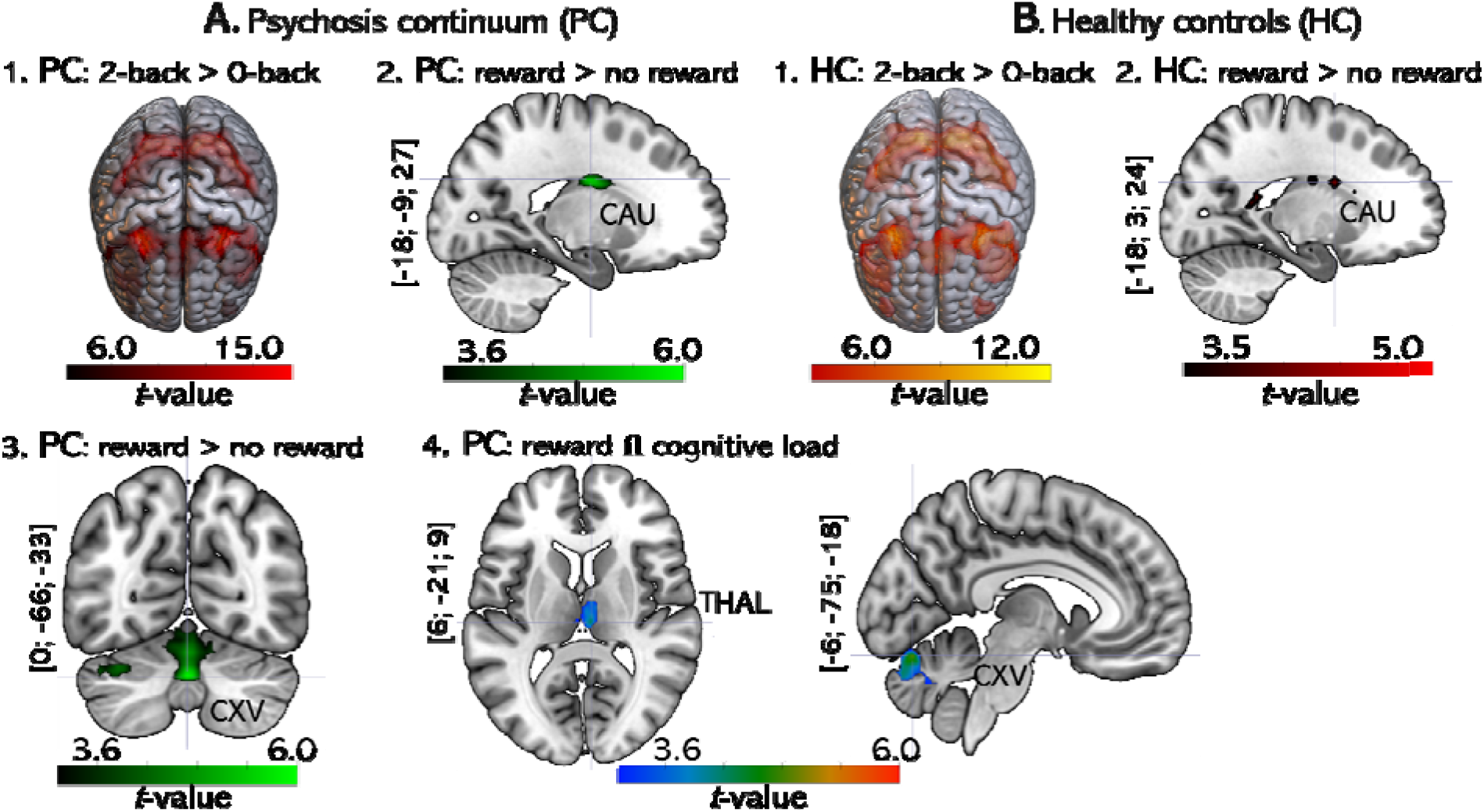
Summary of fMRI results in the psychosis continuum (A) and healthy controls (B). Statistical activations maps are displayed at p-uncorrected<.001 and gray matter masked for illustration purpose; reported activations are corrected for multiple comparisons pFWE<.05 at the cluster level. Abbreviations: CAU=caudate; CXV=vermal cerebellum; THAL=thalamus; PC=psychosis continuum; HC=healthy controls; □ = Conjunction.

#### 3.3.2. Effects of reward

The reward condition (reward>no reward) yielded neural activity in the caudate body ([xyz=(−18; 3; 24), t-value (t)=4.67, cluster extent (ke)=99 voxels, p-value at the cluster level FWE whole-brain corrected for multiple comparisons (p_FWE-CL_)=0.009)] in healthy controls (Figure 2, B2). In the psychosis continuum (Figure 2, A2-3), this contrast (reward>no reward) yielded activity in the vermis [xyz=(0; −66; −33), t=6.00, ke=691 voxels, p_FWE-CL_=10 x −4] and in the left caudate [xyz=(−18; - 9; 27), t=5.91, ke=108, p_FWE-CL_=.001]. Regarding group differences, HC revealed no higher activation during reward condition compared to the psychosis continuum. However, the opposite contrast showed higher cerebellar activity in the psychosis continuum compared to HC [xyz=(−9; −75; −18), t=4.37, ke=170voxels, p_FWE-CL_<0.001].

#### 3.3.3. Conjunction of cognitive load and reward effects

We reasoned that regions computing motivational features and cognitive resource mobilization in the psychosis continuum should activate during both high reward (reward>no reward) and high cognitive load (2-back>0-back). To do so, we first performed a formal conjunction analysis (logical AND) of these contrasts (2-back>0-back □ reward vs no reward). Across the psychosis continuum, the conjunction analysis yielded activity in two clusters in the posterior cerebellum [xyz=(−6; −75; −18); t=4.99 ke=285 voxels, p_FWE-CL_<.001 extending in the vermis [xyz=(0; −60; −33); t=4.20 with a second cerebellar cluster extending more laterally at xyz=(−36; −60; −30); t=4.63, ke=89 voxels, p_FWE-CL_=0.002] and the right thalamus [xyz=(6; −21; 9); t=4.31, ke=61 voxels, p_FWE-CL_=0.015] (see Figure 2, A4). The conjunction analysis for the HC did not reveal any clusters surviving statistical threshold.

#### 3.3.4. Interaction of cognitive load and reward effects

We then examined how reward interacted with cognitive load. Across the psychosis continuum, the interaction analysis [(2-back reward – 2-back no reward)>(0-back reward – 0-back no reward)] showed hemodynamic activity in the left angular gyrus [xyz=(−39; −69;33); t=4.94, ke=125 voxels, p_FWE-CL_<.001] and in the anterior cingulate cortex (ACC) [xyz=(−6; 30; 12); t=4.24, ke=51 voxels, p_FWE-CL_=0.03]. The stronger reward coding under high than low cognitive load is compatible with motivation-related enhancement of cognitive control.

### 3.4. Higher cognitive load relates to negative symptom severity

We examined whether activity in regions involved in either cognitive load or reward processing across the psychosis continuum was differentially associated with clinical scores of negative symptom severity (apathy, diminished expression). Whole brain regression of cognitive load with apathy scores showed a negative association with activity in cortico-striatal-midbrain and cerebellar regions (Figure 3 and Table 2). In particular, the region associated with apathy in the cerebellum overlapped with the region identified in the conjunction analysis at the cerebellar vermal lobules VIII-X [xyz=(−3; −63; −33), t=4.15, p_FWE_=0.002 small volume corrected).

**Figure 3.**
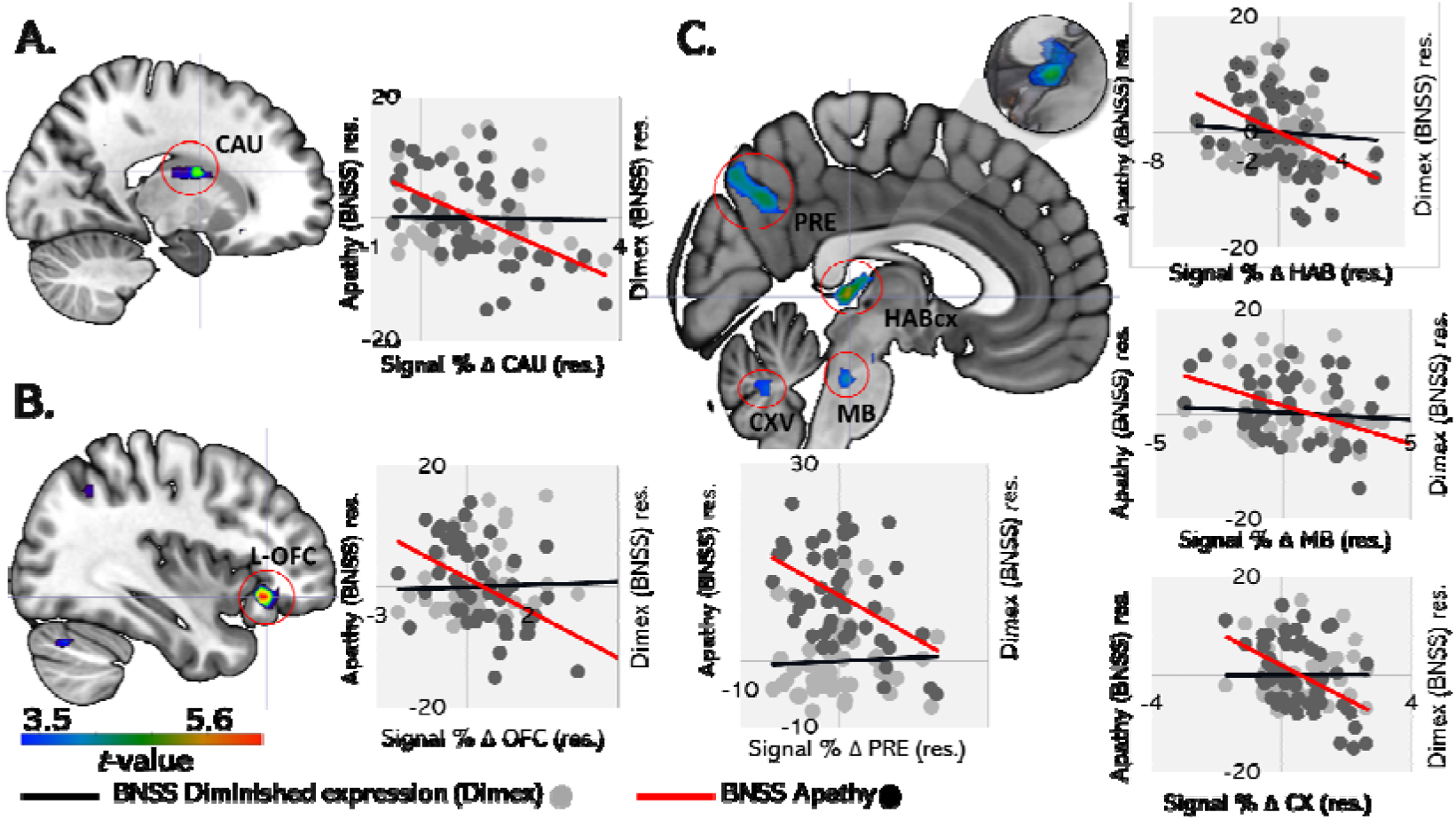
Activation maps of the negative whole-brain regression of cognitive load (2-back>0-back) with apathy scores in the psychosis continuum. Activation maps accompanied by the corresponding scatter plots of partial correlation between brain activity (signal percent change) extracted at cluster peak in the A. nucleus caudate, B. lateral OFC, and C. habenula, precuneus, midbrain and vermis between apathy scores (red line) but not diminished expression scores (black line). Correlation results are corrected for depression scores quantified by the Calgary Depression Scale (CDS). Statistical activations maps displayed at p-uncorrected<.001 and gray matter masked for illustration purpose; reported activations are corrected for multiple comparisons pFWE<.05 at the cluster level. Details on coordinates and statistical significance are found in the main text for the caudate, lateral orbitofrontal cortex, precuneus, habenula, midbrain and vermal cerebellum. Abbreviations: Res.=residuals; OFC=lateral orbitofrontal cortex; CAU=caudate; PRE=precuneus; HAB=habenula; CXV=Vermal cerebellum; MB=Midbrain; BNSS=Brief Negative Symptom Scale; Dimex=diminished expression subscale of BNSS. Apathy=apathy subscale of the BNSS.

Whole brain regression of cognitive load with diminished expression severity did not pass significance threshold. We performed the same regression analysis (apathy and diminished expression) on the reward contrast (reward>no reward) and found no statistically significant results. Moreover, the relationship between apathy and cerebellar activity during cognitive load processing (at xyz=[6; −60; −30]) was significantly stronger than during reward processing (correlation coefficient during cognitive load and reward, respectively −0.48 and 0.18, z=- 3.42, p=6.3 x 10^−4^).

We then asked whether those regions showing an interaction, i.e., stronger reward-related activity under high than low cognitive load, were differentially modulated as a function of negative symptom severity. Whole brain regression of the interaction in the psychosis continuum ([(2-back reward – 2-back no reward)> (0-back reward – 0 back no reward)]) with diminished expression scores revealed a positive correlation at the subgenual ACC [xyz=(0; 21; −9), t=6.11, ke=30, p_FWE_=0.005 corrected at the peak level).

## 4. Discussion

In the current investigation, we aimed to characterize the convergence of reward and cognition processes in the psychosis continuum, and clarify the potential relevance for negative symptoms and in particular for apathy pathophysiology. As expected, higher cognitive load was associated with worse behavioral performance and slower response times, whereas reward improved sensitivity (indexed by *d*’) and accelerated responding. These findings are consistent with the previously established impact of cognitive load and a facilitatory role for reward in performance(Krawczyk, Gazzaley et al. 2007, Braver, Krug et al. 2014, van den Berg, Krebs et al. 2014, Etzel, Cole et al. 2016) and show that reward and cognitive load manipulations were overall effective.

At the neural level, we found that the posterior midline cerebellum and the thalamus were implicated in processing both reward and cognitive load in the psychosis continuum. Reward-related hemodynamic activity in the ACC depended on cognitive load, similar to our previous findings in a chronic schizophrenia cohort employing the same task(Hager, Kirschner et al. 2015). Crucially, reduced hemodynamic activity during cognitive load in the vermal cerebellum, and cortico-striatal-midbrain regions was linked to more severe apathy but not diminished expression in individuals with clinical or subclinical psychosis.

### 4.1. Cerebellar role in flexible modulation of resource allocation

Our results shed light on a key role for the cerebellum in cognitive and reward processes and negative symptom pathophysiology. We show: 1) increased hemodynamic activity in posterior vermal cerebellum during high cognitive load and high reward conditions across the psychosis continuum 2) reduced neural activity during cognitive load in the posterior vermal region in individuals with more severe apathy, but not diminished expression.

A compelling case for cerebellar implication in reward processing comes from animal research where cerebellar climbing fibers – an excitatory input afferent pathway into the cerebellum – signal reward prediction(Heffley and Hull 2019), respond to expected reward size(Larry, Yarkoni et al. 2019) and fire at higher rates for more desired reward outcomes(Heffley, Song et al. 2018, Kostadinov, Beau et al. 2019). Moreover, monosynaptic projections exist from the deep cerebellar nuclei to the VTA(Carta, Chen et al. 2019). Convergent findings in humans indicate that the cerebellum is active during reward anticipation (see meta-analysis of studies employing a monetary incentive delay(Wilson, Colizzi et al. 2018)), while itis also implicated in a range of higher order cognitive processes, in particular WM(E, Chen et al. 2014). Cerebellar activation is cognitive load-dependent, increasing with higher WM load during n-back experimental settings (for review:(Stoodley 2012)). From an anatomical perspective, the cerebellum and the medial frontal cortex are coupled through cerebello – contralateral thalamo – frontal connections(Barch 2014, Palesi, De Rinaldis et al. 2017). Indeed, cerebellar optogenetic stimulation can normalize mediofrontal dysfunction and improve behavior in a WM-attention task(Parker, Kim et al. 2017). Interestingly, in our task we also find contralateral thalamus activation for highly rewarded high-cognitive load trials, which warrants further investigation. Taken together, our results suggest that the cerebellum may be a key hub in the flexible integration of reward and cognitive resources for the modulation of motivated behavior in the psychosis continuum.

### 4.2. Cerebellar role in apathy pathophysiology in the psychosis spectrum

We show for the first time that the midline cerebellum, involved in the reward-cognition interface, also contributes specifically to the pathophysiology of apathy in the psychosis continuum. Indications for cerebellar contributions to global negative symptoms stem from lesion studies in the cerebellum (especially in the posterior lobe), which often manifest as cerebellar cognitive affective syndrome reminiscent of negative symptoms in schizophrenia(Argyropoulos, van Dun et al. 2020). Likewise, the role of the cerebellum in schizophrenia has been recognized in the framework of the Cognitive Dysmetria Theory of schizophrenia(Andreasen, Paradiso et al. 1998) stemming from deficits in cortical-subcortical-cerebellar circuitry. Reduced cerebellar activity was associated with higher global negative symptom severity (indexed by PANSS) in an early psychosis cohort using a Stroop task(Vanes, Mouchlianitis et al. 2019). Cerebellar vermis stimulation was linked to improvement in overall negative symptoms(Demirtas-Tatlidede, Freitas et al. 2010, Tikka, Garg et al. 2015, Garg, Sinha et al. 2016). Recently, dysconnectivity between the rDLPFC and the cerebellar vermis was found as the most significant predictor of negative symptom severity (SANS) and rescuing rDLPFC – cerebellar functional communication was associated with improvement in negative symptom severity(Brady, Gonsalvez et al. 2019). Future well-powered research should tackle the potential of the posterior cerebellum as a therapeutic target for negative symptoms in schizophrenia(Hengyi Cao and Tyrone D. Cannon 2019).

### 4.3. Cortical-striatal-midbrain contributions to apathy in the psychosis continuum

Reduced hemodynamic activity during high cognitive load (2-back>0-back) in cortical areas (lateral OFC, precuneus, lateral parietal cortex (supramarginal gyrus) and middle temporal gyrus), subcortical regions (caudate and habenular complex) and midbrain was associated with more severe apathy in our study.

These results fit with a hypothesized role of parallel thalamocortical and striatal circuits in reward-cognition interface(Cools 2016) where tonic and phasic dopamine release exert dual but distinct functions(Westbrook and Braver 2016). Accordingly, tonic dopaminergic release in the prefrontal cortices, including the lateral OFC(Strauss, Waltz et al. 2014, Westbrook and Braver 2016), favors cognitive *stability* (in other words, the active maintenance of task goals and value representations). Other regions like the lateral parietal cortex (supramarginal gyrus), and to a lesser extent the precuneus and middle temporal gyrus may further stabilize task representations via integration with higher-order motor processes(Tumati, Martens et al. 2019). Phasic release in the striatum facilitates cognitive *flexibility* (the updating of task goals in response to incoming and unexpected information). The caudate belongs to a network of brain regions responsible for the integration of cognitive demands and expected value as the task unfolds(Pessoa and Engelmann 2010, Haber 2016, Doi, Fan et al. 2020). Together with the dopaminergic midbrain, the caudate modulates cognitive resource allocation in a flexible manner in service of goal directed behavior(D’Ardenne, Eshel et al. 2012, Krebs, Boehler et al. 2012, Westbrook and Braver 2016, Hauser, Eldar et al. 2017). The lateral habenula has tight connections with the interpeduncular nucleus of the midbrain and pharmacological lesions of the lateral habenula complex in rats are associated with decreased willingness to exert effort for high-reward high-effort trials(Sevigny, Bryant et al. 2021). Lesions of the habenula manifest with negative and cognitive schizophrenia-like behavior in rats via downregulation of the D1R expression in the medial PFC(Li, Yang et al. 2019). Future studies should investigate the role of the human habenular complex in negative symptoms of schizophrenia(Shepard, Holcomb et al. 2006).

### 4.4. Anterior cingulate – a role in diminished expression pathophysiology even in early stages of psychosis

We extend our previous findings in the chronic schizophrenia population (Hager, Kirschner et al. 2015) to earlier stages of the psychosis spectrum. Indeed, in the current study, we found that hemodynamic activity in the ACC was associated to stronger reward modulation of high vs. low cognitive load (interaction analysis), similar to our previous findings in a chronic schizophrenia cohort employing the same task(Hager, Kirschner et al. 2015). This result suggests that even at early stages of psychosis, the ACC is recruited to orchestrate and prioritize cognitive resource capacities when rewards are at stake, an observation which is in line with the role of the ACC in controlling cognitive resource distribution(Holroyd and Yeung 2012). However, in contrast to our findings in the chronic schizophrenia cohort, we found that ACC activity was positively (not negatively(Hager, Kirschner et al. 2015)) associated to diminished expression but not apathy severity. Together with findings of preserved frontal capacity(Siever and Davis 2004) and neuroanatomy in schizotypy(Kirschner, Hodzic-Santor et al. 2021), the present results suggest that in subclinical and minimally-treated stages of the psychosis continuum, more severe negative symptoms (diminished expression) are linked to an increased compensatory recruitment of cognitive capacities, operated by the ACC. This mechanism may saturate in later more chronic stages of psychosis, even though not as strongly as to impair task performance in chronic patients(Hager, Kirschner et al. 2015). More work is needed to better understand compensatory processes in the psychosis spectrum. Whereas saturation of compensatory mechanisms could be one explanation, our divergent results in early vs. chronic phases of psychosis (e.g. positive correlation of ACC activity with diminished expression in early stages but negative correlation in chronic schizophrenia) may translate distinct clinical trajectories. Thus, it remains to be clarified whether observations in schizophrenia result from the breakdown of the same mechanism as in early psychosis individuals or whether it indicates a different “schizophrenia-specific” mechanism. Longitudinal data from first-episode schizophrenia cohorts and a matched group of healthy schizotypy would be important to test the hypothesis of distinct trajectories.

## 5. Strengths, limitations and opportunities for future research

Our study cohort has strength and limitations. A main strength is that in the current study we excluded obvious sources of secondary negative symptoms (e.g., due to depression, psychosis, medication and sedation). In addition, considering that chronic antipsychotic medication exposure can modulate or change the relationship between negative symptoms and neural activity, another strength is the inclusion of unmedicated and minimally treated individuals allowing us to examine changes in early psychosis stages. Finally, the present study provides evidence for specific functional neural signatures related to diminished expression and to apathy in the psychosis continuum providing further support for a dimension-specific approach to negative symptoms of schizophrenia.

The main limitation in our study is the relatively modest sample size. Larger sample sizes would be necessary to better characterize the correlates of specific symptom domains for the individual subgroups.

Another limitation relates to our conceptual framework: in the current study, we applied the concept of the psychosis continuum that assumes that schizotypy represents early subclinical stages of psychosis, based on evidence showing shared neuroimaging and genetic features with psychotic disorders(DeRosse and Karlsgodt 2015) in addition to a role for dopaminergic transmission even at this stage(Mohr and Ettinger 2014). With respect to the schizotypy participants, considering most of these individuals will not develop psychosis(Debbané, Eliez et al. 2015), our crosssectional design does not allow us to differentiate between healthy schizotypy with high subclinical negative symptoms and schizotypy as at-risk stage for psychosis.

Our study opens interesting future research opportunities in particular to directly and causally investigate the role of the cerebellum, and the cortico-cerebellar circuitry in general, in apathy pathophysiology affecting the psychosis spectrum. Moreover, the inclusion of additional affective disorders would be required to clarify whether deficits in cortical-striatal-midbrain-cerebellar circuitry are specific for the psychosis continuum or represent a transdiagnostic signature of apathy.

## 6. Conclusions

We investigated the behavioral and neural convergence of cognitive load and reward processes in the psychosis continuum and their contribution to amotivational negative symptoms. We show that hypoactivity in the cerebellar vermis in addition to several reward-cognition nodes in cortical-striatal-midbrain-cerebellar circuitry relates to apathy severity in the psychosis continuum. Moreover, our findings suggest that a clinically-relevant cerebellar mechanism may underpin the flexible modulation of behavior by reward and cognitive resource allocation for effective goal pursuit in the psychosis continuum.

## Acknowledgment

We are grateful to Dr. Philipp Staempfli for his excellent MRI support.

## Contributors

S. Kaiser and P. Tobler designed the study. J. Brakowski and M. Kirschner conducted the study. I. Bègue conducted the analyses and wrote the first draft of the manuscript. I.Bègue, J. Brakowski, E. Seifritz, A.Dagher, P. Tobler, M. Kirschner and S. Kaiser contributed to and have approved the final manuscript.

