## Supplementary material for "Cerebellar and cortico-striatal-midbrain contributions to reward-cognition processes and apathy within the psychosis continuum"

Indrit Bègue (indrit dot begue (at) unige.ch)

^1^Division of Adult Psychiatry, Department of Psychiatry, Geneva University Hospitals, Switzerland

^*^M.K. and S.K. contributed equally to this work

**S1. Participants**

Patients with FEP were recruited from outpatient and inpatient units of the Psychiatric Hospital of the University of Zurich with a clinical diagnosis of brief psychotic disorder or schizophreniform disorder confirmed in a structured Mini-International Neuropsychiatric Interview (M.I.N.I)(Sheehan, Lecrubier et al. 1997). We excluded participants with any other DSM-IV axis I disorder (in particular current substance use disorder and substance-induced psychotic disorder). Moreover, we excluded patients with major causes of secondary NS, i.e. with major depressive episode, significant extrapyramidal side effects (defined as a total score higher than 2 on the Modified Simpson–Angus Scale(Simpson and Angus 1970)), florid psychotic symptoms (defined as any positive subscale item score four or higher measured with the Positive and Negative Syndrome Scale (PANSS)(Kay, Opler et al. 1987)) or excessive sedation (lorazepam more than 1 mg/d). Twenty patients received a constant dose of atypical antipsychotic with no change in medication for at least 14 days prior to testing, while three patients were not on medication. Individuals with schizotypal traits were recruited using an online form of the Schizotypal Personality Questionnaire (SPQ)(Raine 1991). 956 participants completed the questionnaire (mean score=16.66, SD=11.34). Individuals with the highest SPQ total scores (upper 10% of the SPQ total score) were invited to participate in the study. Exclusion criteria were any current or past Axis I disorder confirmed with the M.I.N.I(Sheehan, Lecrubier et al. 1997) as well as use of psychopharmacological drugs. The local Ethics committee of the Canton Zurich, Switzerland approved the study, and all participants gave written informed consent.

All individuals of the psychosis continuum underwent a detailed psychopathological assessment. To assess the two symptom dimensions of apathy and diminished expression, we used the validated German version of the Brief Negative Symptoms Scale (BNSS)(Strauss, Hong et al. 2012, Bischof, Obermann et al. 2016). Additionally, we employed the Positive and Negative Syndrome Scale (PANSS)(Kay, Opler et al. 1987), the Calgary Depression Scale for Schizophrenia (CDS)(Addington, Addington et al. 1993), the Global Assessment of Functioning scale (GAF)(Association 2000) and the Personal and Social Performance Scale (PSP)(Schaub and Juckel 2011). To characterize neuropsychological functioning the following cognitive domains were tested in the complete sample: verbal learning (Auditory Verbal Learning Memory Test)(Helmstaedter and Durwen 1990), verbal and visual short-term working memory (Digit span, DS)(Stieglitz 2000), Corsi block-tapping (CBT)(Kessels, Van Zandvoort et al. 2000), processing speed (Digit-Symbol Coding)(Molz, Schulze et al. 2010), planning (Tower of London)(Shallice 1982), semantic and phonetic fluency (animal naming, s-words)(Delis DC 2001) and premorbid intelligence (Multiple Word Test Intelligence Quotient, MWT IQ)(Lehrl, Triebig et al. 1995).

| **Table S1**. Summary of the relationship of clinical and neuropsychological data with negative symptoms. | | | | |
| --- | --- | --- | --- | --- |
|  | ***N*** | **Spearman rank order correlation coefficient** | **t(N-2)** | **p-value (uncorrected)** |
| ***Psychosis (PANSS positive factor)*** |  |  |  |  |
| x BNSS Dimex | 50 | -0.19 | -1.41 | 0.16 |
| x BNSS Apathy | 50 | 0.30 | 2.18 | 0.03 |
| ***Medication dosage (Chlorpromazine equivalents)*** |  |  |  |  |
| x BNSS Dimex | 23 | 0.23 | 1.09 | 0.29 |
| x BNSS Apathy | 23 | -0.13 | -0.60 | 0.55 |
| ***General functioning (GAF)*** |  |  |  |  |
| x BNSS Dimex | 49 | -0.19 | -1.37 | 0.18 |
| x BNSS Apathy ***** | 49 | -0.69 | -6.47 | .05 x 10 -6 |
| ***Psychosocial functioning (PSP)*** |  |  |  |  |
| x BNSS Dimex | 48 | -0.25 | -1.77 | 0.08 |
| x BNSS Apathy ***** | 48 | -0.70 | -6.65 | 0.03 x 10 -6 |
| ***Depression (CDS)*** |  |  |  |  |
| x BNSS Dimex | 50 | .057 | 0.39 | 0.69 |
| x BNSS Apathy ***** | 50 | 0.47 | 3.69 | 0.0005 |
| ***Intelligence (MWTQ)*** |  |  |  |  |
| x BNSS Dimex ***** | 49 | -0.46 | -3.59 | 0.0008 |
| x BNSS Apathy | 49 | -0.18 | -1.27 | 0.21 |
| ***Auditory verbal learning memory (VLMT)*** |  |  |  |  |
| x BNSS Dimex | 50 | -0.18 | -1.29 | 0.20 |
| x BNSS Apathy | 50 | -0.24 | -1.75 | 0.09 |
| ***Verbal and visual short-term working memory (Digit Span, DS)*** |  |  |  |  |
| *DS forward* |  |  |  |  |
| x BNSS Dimex | 50 | -0.28 | -2.06 | 0.04 |
| x BNSS Apathy | 50 | -0.31 | -2.30 | 0.02 |
| *DS backward* |  |  |  |  |
| x BNSS Dimex | 50 | -0.19 | -1.39 | 0.16 |
| x BNSS Apathy | 50 | -0.13 | -0.89 | 0.37 |
| *DS total score* |  |  |  |  |
| x BNSS Dimex | 50 | -0.25 | -1.79 | 0.08 |
| x BNSS Apathy | 50 | -0.23 | -1.65 | 0.10 |
| ***Processing speed (Digit-Symbol Coding, DSC)*** |  |  |  |  |
| x BNSS Dimex | 50 | -0.20 | -1.42 | 0.16 |
| x BNSS Apathy | 50 | -0.24 | -1.74 | 0.08 |
| ***Short memory task (Corsi block-tapping test, CBT)*** |  |  |  |  |
| *CBT forward* |  |  |  |  |
| x BNSS Dimex | 50 | 0.07 | 0.50 | 0.62 |
| x BNSS Apathy | 50 | -0.07 | -0.50 | 0.61 |
| *CBT backward* |  |  |  |  |
| x BNSS Dimex | 50 | -0.06 | -0.39 | 0.69 |
| x BNSS Apathy | 50 | 0.01 | 0.048 | 0.96 |
| *CBT total* |  |  |  |  |
| x BNSS Dimex | 50 | -0.03 | -0.194 | 0.85 |
| x BNSS Apathy | 50 | -0.03 | -0.20 | 0.84 |
| ***Executive functioning (Tower of London, TOL)*** |  |  |  |  |
| x BNSS Dimex | 50 | -0.29 | -2.11 | 0.04 |
| x BNSS Apathy | 50 | -0.27 | -1.91 | 0.06 |
| ***Verbal fluency (letter "s")*** |  |  |  |  |
| x BNSS Dimex | 50 | -0.38 | -2.85 | 0.006 |
| x BNSS Apathy | 50 | -0.25 | -1.81 | 0.07 |
| ***Verbal fluency ('animals')*** |  |  |  |  |
| x BNSS Dimex | 50 | -0.33 | -2.40 | 0.02 |
| x BNSS Apathy | 50 | -0.24 | -1.70 | 0.09 |
| *Abbreviations*: x=correlation. *BNSS=Brief Negative Symptom Scale. Dimex=Diminished expression. Significance threshold set at p=.001 6 (corresponding to Bonferroni correction for multiple comparisons for 32 tests). Significant correlation marked with* ******* | | | | |

| **Table S2**. Summary of brain activation during cognitive load (2-back > 0-back) | | | | | | |
| --- | --- | --- | --- | --- | --- | --- |
|  | |  |  | **MNI Coordinates** | | |
| Region Label | | Cluster extent | t-value | x | y | z |
| *Psychosis continuum* | | | | | | |
| L | Supramarginal gyrus | 6399 | 17.61 | -36 | -42 | 39 |
| R | Cerebellum (Crus 1) |  | 15.96 | 30 | -63 | -30 |
| L | Precuneus |  | 15.39 | -9 | -69 | 54 |
| L | Middle Frontal Gyrus | 5703 | 16.19 | -24 | 0 | 57 |
| R | Middle Frontal Gyrus |  | 15.75 | 27 | 6 | 57 |
| L | Posterior-Medial Frontal Cortex | | 14.68 | -3 | 9 | 51 |
| *Healthy controls* | | | | | | |
| L | Precuneus | 3467 | 12.69 | -9 | -66 | 54 |
| L | Superior Parietal Lobule |  | 12.57 | -33 | -45 | 42 |
| R | Precuneus |  | 12.49 | 12 | -66 | 54 |
| L | Middle Frontal Gyrus | 4726 | 12.34 | -27 | 3 | 51 |
| R | Middle Frontal Gyrus |  | 11.47 | 27 | 9 | 51 |
| L | Posterior-Medial Frontal |  | 11.41 | -6 | 12 | 51 |
| R | Cerebellum (Crus 1) | 868 | 12.01 | 33 | -66 | -30 |
| L | Cerebellum (Crus 1) |  | 10.42 | -36 | -66 | -30 |
| R | Cerebellum (VIII) |  | 8.20 | 9 | -75 | -27 |
| R | Inferior frontal gyrus pars orbitofrontalis | 116 | 7.05 | 33 | 30 | 0 |
| *Table shows all local maxima separated by more than 20 mm. Regions were automatically labeled using the AAL3 atlas and Neuromophometrics in SPM12. x, y, and z =Montreal Neurological Institute (MNI) coordinates in the left-right, anterior-posterior, and inferior-superior dimensions, respectively. Thresholding at t > 3.1273; p<.001 0; df=222; minimum extent=30.* | | | | | | |

| **Table S3**: Group differences for the cognitive load contrast (2back vs 0back) | | | | | | |
| --- | --- | --- | --- | --- | --- | --- |
|  |  |  |  | MNI Coordinates | | |
|  | Region Label | Extent | t-value | x | y | z |
| PC > HC | | | | | | |
|  | L Inferior Parietal Lobule | 1767 | 8.245 | -30 | -54 | 51 |
|  | R Superior Frontal Gyrus | 1981 | 7.996 | 27 | 0 | 51 |
|  | R Cerebellum (VI) | 327 | 6.983 | 27 | -63 | -30 |
|  | R IFG (p. Orbitalis) | 48 | 4.659 | 33 | 24 | -3 |
|  | R Middle Frontal Gyrus | 51 | 4.581 | 36 | 30 | 39 |
| HC > PC | | | | | | |
|  | L Insula Lobe | 244 | -6.348 | -39 | -15 | 18 |
|  | R Rolandic Operculum | 404 | -5.515 | 48 | -12 | 18 |
|  | R Superior Medial Gyrus | 255 | -5.199 | 3 | 60 | 6 |
| *Table shows all local maxima separated by more than 20 mm for group differences for the contrast 2back vs 0 back. Regions were automatically labeled using the AnatomyToolbox atlas. x, y, and z =Montreal Neurological Institute (MNI) coordinates in the left-right, anterior-posterior, and inferior-superior dimensions, respectively. Thresholding t > 3.1273; p < 0.0010; df = 222. All activations surviving pfwe corrected at the cluster level < 0.05. Abbreviations PC = psychosis continuum; HC = Healthy controls* | | | | | | |

Addington, D., J. Addington and E. Maticka-Tyndale (1993). "Assessing depression in schizophrenia: the Calgary Depression Scale." The British journal of psychiatry **163**(S22): 39-44.

Association, A. P. (2000). "Global assessment of functioning scale." Diagnostic and statistical manual of mental disorders, 4th edition, text revision. Washington, DC: American Psychiatric Association **34**: 39.

Bischof, M., C. Obermann, M. N. Hartmann, O. M. Hager, M. Kirschner, A. Kluge, G. P. Strauss and S. Kaiser (2016). "The brief negative symptom scale: validation of the German translation and convergent validity with self-rated anhedonia and observer-rated apathy." BMC Psychiatry **16**(1): 415.

Delis DC, K. E., Kramer J. Delis Kaplan (2001). "Executive Function System. ." The Psychological Corporation; 2001: San Antonio, TX, 2001.

Helmstaedter, C. and H. Durwen (1990). "VLMT: Verbaler Lern-und Merkfähigkeitstest: Ein praktikables und differenziertes Instrumentarium zur Prüfung der verbalen Gedächtnisleistungen." Schweizer Archiv für Neurologie, Neurochirurgie und Psychiatrie.

Kay, S., A. Opler, A. Fiszbein, P. Ramirez and L. White (1987). "The Positive and Négative Syndrome Scale for schizophrénia." Schizophr Bull **3**: 26-76.

Kessels, R. P., M. J. Van Zandvoort, A. Postma, L. J. Kappelle and E. H. De Haan (2000). "The Corsi block-tapping task: standardization and normative data." Applied neuropsychology **7**(4): 252-258.

Lehrl, S., G. Triebig and B. Fischer (1995). "Multiple choice vocabulary test MWT as a valid and short test to estimate premorbid intelligence." Acta Neurol Scand **91**(5): 335-345.

Molz, C., R. Schulze, U. Schroeders and O. Wilhelm (2010). "Wechsler Intelligenztest für Erwachsene WIE. Deutschsprachige Bearbeitung und Adaptation des WAIS-III von David Wechsler." Psychol. Rundsch **61**: 229-230.

Raine, A. (1991). "The SPQ: A Scale for the Assessment of Schizotypal Personality Based on DSM-III-R Criteria." Schizophrenia Bulletin **17**(4): 555-564.

Schaub, D. and G. Juckel (2011). "PSP-Skala—Deutsche version der Personal and Social Performance Scale: Validiertes messinstrument zur erfassung des psychosozialen funktionsniveaus in der schizophrenietherapie." Der Nervenarzt.

Shallice, T. (1982). "Specific impairments of planning." Philosophical Transactions of the Royal Society of London. B, Biological Sciences **298**(1089): 199-209.

Sheehan, D., Y. Lecrubier, K. H. Sheehan, J. Janavs, E. Weiller, A. Keskiner, J. Schinka, E. Knapp, M. Sheehan and G. Dunbar (1997). "The validity of the Mini International Neuropsychiatric Interview (MINI) according to the SCID-P and its reliability." European psychiatry **12**(5): 232-241.

Simpson, G. and J. Angus (1970). "A rating scale for extrapyramidal side effects." Acta Psychiatrica Scandinavica **45**(S212): 11-19.

Stieglitz, R. (2000). "WMS-R. Wechsler Gedächtnistest-revidierte Fassung." Z. Für Klin. Psychol. Psychother **29**: 307-308.

Strauss, G. P., L. E. Hong, J. M. Gold, R. W. Buchanan, R. P. McMahon, W. R. Keller, B. A. Fischer, L. T. Catalano, A. J. Culbreth and W. T. Carpenter (2012). "Factor structure of the brief negative symptom scale." Schizophrenia research **142**(1-3): 96-98.
